# Reduced bushy cell firing limits binaural coding after cochlear synaptopathy

**DOI:** 10.64898/2026.11.21.768331

**Authors:** Go Ashida, Daniel J. Tollin

## Abstract

Listeners can experience difficulties with spatial hearing and speech understanding in complex acoustic environments despite near-normal hearing thresholds. Cochlear synaptopathy, which reduces auditory nerve input without elevating thresholds, is one candidate mechanism, but how this peripheral deafferentation disrupts binaural processing remains unclear. We adapted a physiologically constrained computational model incorporating the auditory periphery, spherical and globular bushy cells of the cochlear nucleus and the lateral superior olive (LSO) to examine the separate and combined effects of reduced afferent input and degraded spike timing. Synaptopathy was modeled as loss of auditory nerve inputs to bushy cells, whereas central myelin dysfunction was modeled as increased temporal jitter in bushy cell outputs. Loss of auditory nerve input substantially reduced bushy cell firing rates without degrading phase locking. Despite this preserved temporal precision, binaural tuning in the LSO declined disproportionately, where moderate afferent loss produced substantial reductions in sensitivity to interaural envelope phase differences. In contrast, temporal jitter preserved bushy cell firing rates but degraded phase locking and binaural tuning. Selective loss of input to excitatory versus inhibitory pathways produced opposite changes in overall LSO firing rate, yet both reduced binaural interaction. These results identify two mechanistically distinct routes to impaired binaural coding: too few precisely timed spikes, and a sufficient number of spikes arriving with inadequate temporal precision. These results predict that correcting myelin-related timing deficits alone would leave substantial binaural coding impairment when cochlear deafferentation persists, motivating interventions that address both diminished afferent drive and impaired transmission.

## Introduction

A sound can remain audible even when the neural representation necessary to locate or separate it from competing sounds has deteriorated. Listeners with clinically normal thresholds differ in their ability to understand spatially separated competing talkers, with performance associated with subcortical temporal coding [1]. Cochlear synaptopathy, the loss of synapses between inner hair cells and auditory nerve (AN) fibers, offers one possible explanation. Noise exposure and aging can reduce cochlear innervation without commensurate threshold elevation [2, 3], although associations between proposed measures of human synaptopathy and listening difficulties remain inconsistent [4, 5]. Understanding its functional consequences requires determining how surviving input is transformed by central auditory circuits.

An important distinction is between spike precision and availability for downstream computation. Bushy cells in the cochlear nucleus can sharpen phase locking, the alignment of spikes with a recurring sound phase, relative to their AN inputs [6, 7]. This temporal sharpening was demonstrated to improve binaural envelope coding in a recent circuit model [8]. Precise phase locking can persist while firing rate decreases. If downstream computation requires several inputs to arrive within a short interval, surviving spikes may be accurately timed but too sparse to support it [9]. Synaptopathy could therefore impair binaural processing despite preserved monaural temporal precision and overall audibility.

The lateral superior olive (LSO) provides a circuit to test this possibility. Spherical bushy cells (SBCs) supply ipsilateral excitation, whereas globular bushy cells (GBCs) supply contralaterally driven inhibition through the medial nucleus of the trapezoid body (MNTB) (Fig. 1A) [10, 11]. Their differing synaptic input arrangements could confer different vulnerabilities to deafferentation. These pathways support sensitivity to interaural level differences (ILDs) and interaural time differences (ITDs) in amplitude-modulated (AM) sound envelopes [12]. Modeling [13] and recordings [14] show that LSO responses depend on excitatory coincidences and their timing relative to inhibition, allowing input loss to be examined across successive coincidence sensitive stages, the bushy cells and the LSO.

**Fig. 1.**
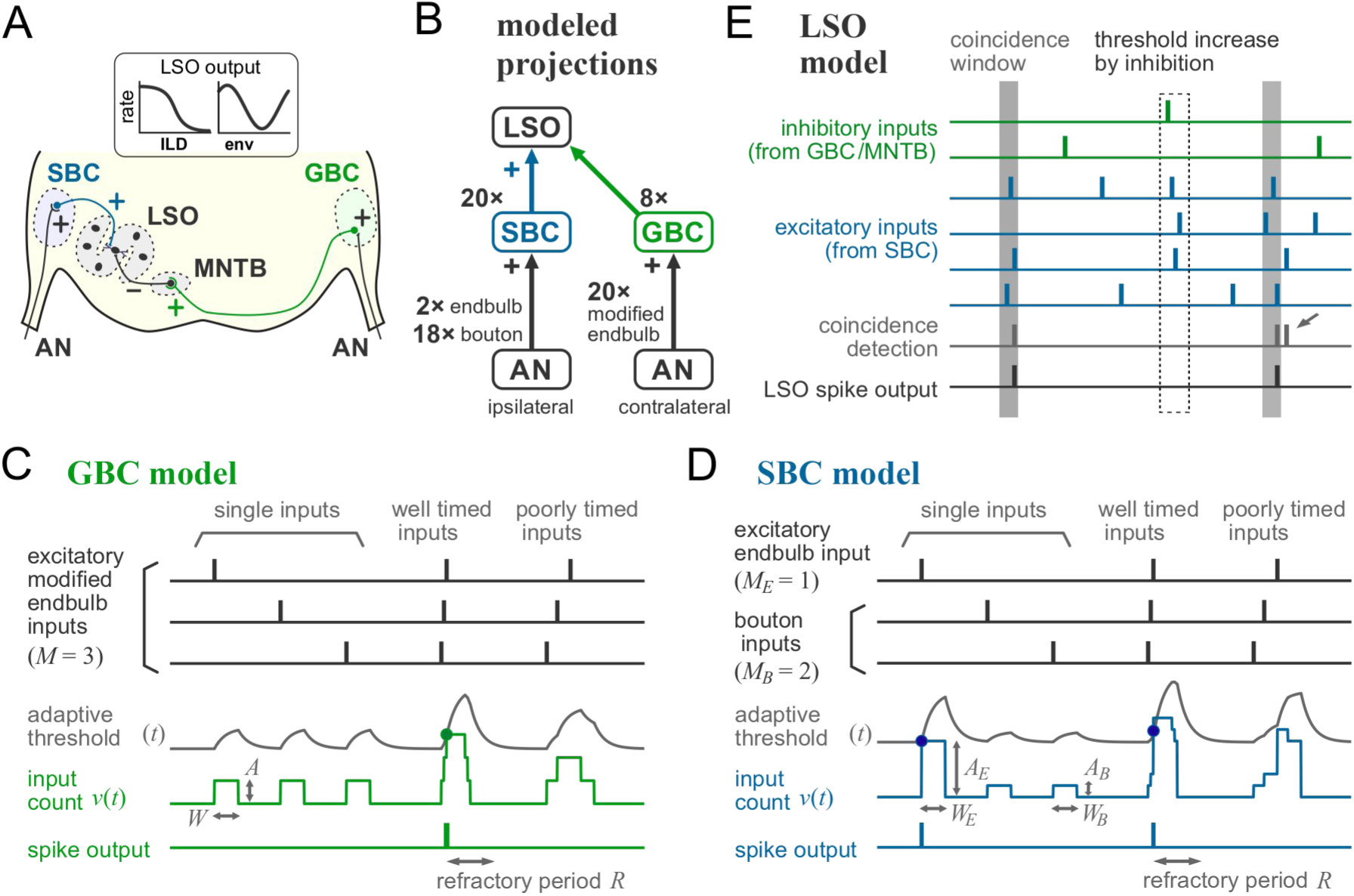
| Architecture of the auditory nerve and brainstem circuit. **(A)** LSO circuit. AN activity drives ipsilateral SBCs and contralateral GBCs. SBCs excite the LSO, while GBCs drive inhibition through the MNTB. Plus and minus signs indicate excitation and inhibition, respectively. The inset shows schematic rate tuning to ILD and envelope ITD. **(B)** Intact convergence: 20 AN inputs per bushy cell, 20 excitatory SBC inputs and eight GBC-driven inhibitory inputs per LSO neuron. MNTB transmission is represented by sign inversion. **(C)** LSO operation. Light blue and light green events indicate excitatory and inhibitory inputs, respectively. Gray bands mark coincidence windows, and the dashed box marks threshold increase caused by inhibition. Lower traces indicate detected coincidences and output spikes. Refractoriness prevents an additional output at the arrow. **(D)** GBC operation. An input of amplitude *A* and duration *W* contributes to the summed spike count *v*(*t*). Coincident inputs that cross the adaptive threshold *θ*(*t*) generate an output followed by a refractory period *R*. Three inputs are shown schematically; simulations used 20. **(E)** SBC operation. Endbulb and bouton responses have amplitudes *A*_E_ and *A*_B_ and durations *W*_E_ and *W*_B_, respectively. One endbulb and two bouton inputs are shown schematically; simulations used two endbulbs and 18 boutons. Dots mark threshold crossings. **AN**, auditory nerve; **SBC**, spherical bushy cell; **GBC**, globular bushy cell; **LSO**, lateral superior olive; **MNTB**, medial nucleus of the trapezoid body; **ILD**, interaural level difference; **ITD**, interaural time difference. Model components are based on refs. [8, 9, 11, 13].

Spike transmission timing provides a second vulnerability. Axonal geometry and myelination help regulate action potential arrival times [15], and myelin defects increase timing variability and conduction failures [16]. Auditory deprivation reduces myelin thickness in adult trapezoid-body fibers [17] and produces axon-specific myelination differences during development [18]. More broadly, neuronal activity can regulate oligodendrocyte generation and axonal myelination [19, 20]. These findings raise a conditional hypothesis. If synaptopathy persistently reduces bushy-cell output, altered activity-dependent myelin regulation could subsequently compromise spike transmission along surviving axons. Input loss and timing disruption might then be successive consequences of synaptopathy.

Previous models demonstrated impaired low-frequency ITD coding in the medial superior olive (MSO) after cochlear synaptopathy or AN myelin defects [21], and sensitivity of MSO tuning to central axonal myelination [22]. Other simulations found relatively robust LSO tuning after direct inhibitory input loss, particularly with compensatory synaptic strengthening [23]. How the distinct SBC and GBC afferent arrangements transform AN input loss into LSO envelope coding deficits, and how these deficits interact with impaired transmission timing, remains unresolved. A related therapeutic question is whether correcting transmission timing could restore binaural processing when persistent deafferentation due to synaptopathy continues to limit the available inputs.

Here we combine an auditory periphery model [24] with physiologically constrained bushy cell and LSO models [8, 9, 11, 13]. Removing AN inputs represents functional deafferentation, while adding jitter to bushy cell outputs isolates temporal dispersion. We assess firing, phase locking, binaural tuning and neural discriminability after separate, combined and pathway-selective perturbations. Afferent loss disproportionately weakens binaural coding despite largely preserved bushy cell phase locking, and timing dispersion compounds the impairment. These results identify the supply of well-timed spikes as a determinant of binaural coding and motivate tests of secondary, activity-dependent myelin dysfunction.

## Results

### A multistage auditory circuit separates input availability from transmission timing

We connected a model of the auditory periphery [24] to SBC [8] and GBC [9] stages in the cochlear nucleus and a downstream binaural LSO neuron model [11, 13] to separate afferent loss from temporal dispersion (Fig. 1). Each bushy cell received 20 AN inputs. GBC inputs were equally weighted and subthreshold, whereas SBCs received two large endbulb and 18 small bouton inputs (Fig. 1B). Both models generated spikes when summed input reached an adaptive threshold outside the refractory period (Fig. 1C,D). The LSO received 20 excitatory SBC inputs and eight inhibitory inputs through GBCs and the MNTB, with MNTB transmission represented by sign inversion. Excitation accumulated within a coincidence window, while inhibition temporarily raised the firing threshold (Fig. 1E) [13].

We removed AN inputs to bushy cells to model functional deafferentation and added jitter after bushy cell spike generation to impair timing without deleting spikes. Perturbations affected both bushy cell pathways. For AM envelope frequencies of 150, 300 and 450 Hz, we measured bushy cell rate and vector strength (VS, a measure of phase locking), and LSO tuning and neural discriminability to envelope interaural phase difference (IPD) or ILD.

### Afferent loss reduces bushy-cell firing while largely preserving phase locking

We first compared rate-level functions and envelope phase locking with intact AN input and after 25% or 50% input loss (Fig. 2). In the intact model, firing rates increased at low-to-moderate sound levels and then approached a plateau. GBC rates decreased shallowly at the highest levels for the lower modulation frequencies (Fig. 2A1-A3, blue), whereas SBC rates remained comparatively stable after saturation (Fig. 2B1-B3, blue). SBC firing exceeded GBC firing across the illustrated conditions, but GBCs showed stronger envelope phase locking (Fig. 2C1-C3,D1-D3). This greater temporal sharpening reflected coincidence detection of multiple subthreshold inputs [8, 9].

**Fig. 2.**
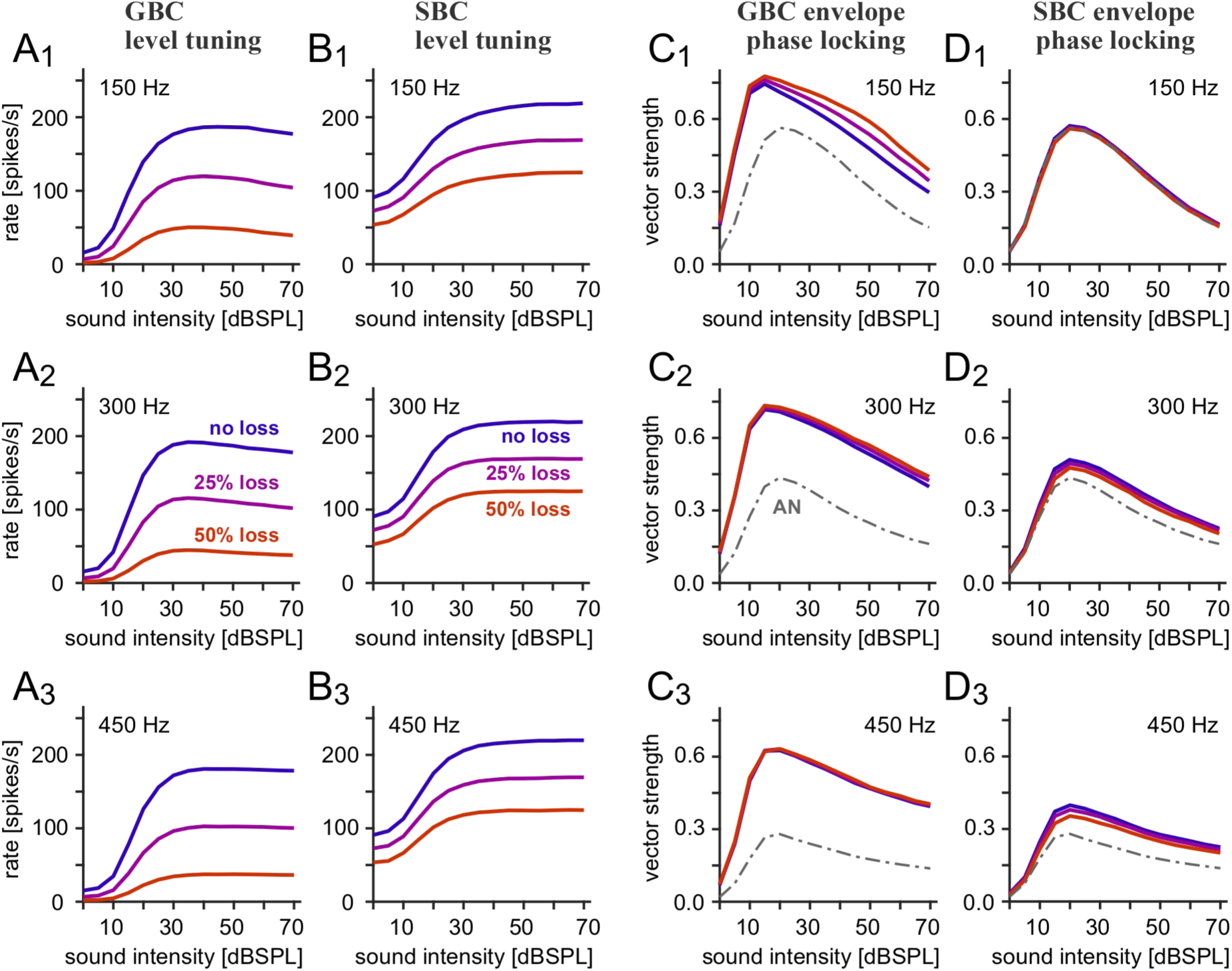
| Bushy-cell firing rates and phase locking after auditory nerve input loss. Responses to AM sounds are plotted against sound level. Rows 1–3 show envelope frequencies of 150, 300 and 450 Hz. **(A_1_–A_3_)** GBC firing rates. **(B_1_–B_3_)** SBC firing rates. **(C_1_–C_3_)** GBC envelope phase locking quantified by VS. **(D_1_–D_3_)** SBC envelope phase locking quantified by VS. Blue, purple, and orange curves indicate intact input, 25% input loss and 50% input loss, respectively. Gray dash-dotted curves show the AN reference where visible. Loss refers to functional AN inputs to the indicated bushy-cell population; no additional output jitter was applied.

Input loss reduced firing in both populations at all three envelope frequencies, with larger reductions in GBCs (purple and orange curves in Fig. 2A1-A3,B1-B3). Increasing sound level did not recover the lost GBC response. SBCs retained a larger fraction of their output, although their rate-level functions also shifted downward. VS remained comparatively stable, preserving its rise at low-to-moderate levels and decline at higher levels (Fig. 2C1-C3,D1-D3). GBC phase locking increased modestly under some conditions, particularly at 150 Hz. SBC phase locking changed little at 150 Hz and decreased slightly at higher envelope frequencies. Deafferentation therefore primarily reduced spike availability while largely preserving phase locking.

### Binaural tuning declines disproportionately despite preserved phase locking precision

We next traced progressive AN input loss through both bushy cell pathways to the LSO at 35 dB SPL, where binaural IPD tuning was pronounced [8]. SBC firing declined approximately in proportion to input loss, whereas GBC output declined more steeply (Fig. 3A1-A3). At 50% loss, GBCs retained approximately one-quarter of their intact firing rate and SBCs retained about a half. This difference followed from their input configurations. Reducing convergence sharply decreased the probability that multiple subthreshold inputs coincided in GBCs, while each SBC endbulb input was normally sufficient to generate an output spike [8]. Across the same range of loss, GBC VS was maintained or increased slightly, and SBC VS remained nearly constant or decreased modestly (Fig. 3B1-B3).

**Fig. 3.**
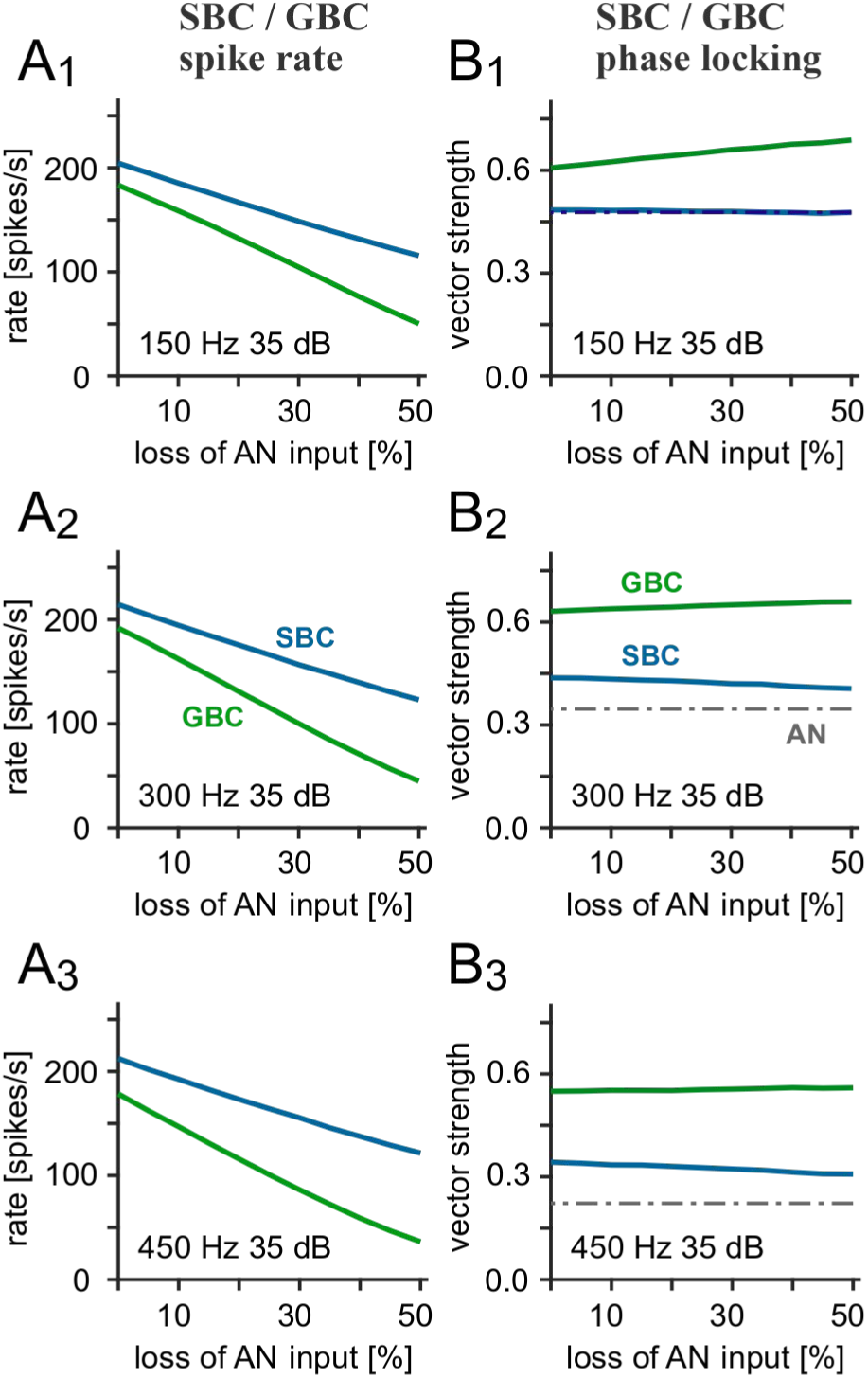
| Bushy-cell responses across degrees of input loss. Responses were simulated at 35 dB SPL with matched fractional AN input loss in both bushy-cell pathways and no additional jitter. Rows 1–3 show envelope frequencies of 150, 300 and 450 Hz. **(A_1_–A_3_)** Bushy-cell rates versus input loss. Light blue and light green indicate SBCs and GBCs, respectively. **(B_1_–B_3_)** Corresponding VS values. Gray dash-dotted lines indicate the AN reference.

LSO tuning to IPD nevertheless deteriorated markedly. Intact responses at 150 and 300 Hz showed a broad maximum at negative envelope IPDs and a trough near zero IPD (Fig. 4A1,A2, blue). The intact 450-Hz response was also phase dependent but had a lower peak and shallower tuning (Fig. 4A3, blue). Removing 25% of AN inputs (purple curves) reduced tuning depth by approximately half at 150 Hz and by more than half at 300 and 450 Hz. Removing 50% (red curves) left weak residual modulation at 150 Hz and suppressed most phase-dependent firing at higher frequencies. The maximum spike rate fell while the minimum remained low (Fig. 4B1-B3), producing fractional losses of tuning depth that exceeded AN input loss over much of the tested range.

**Fig. 4.**
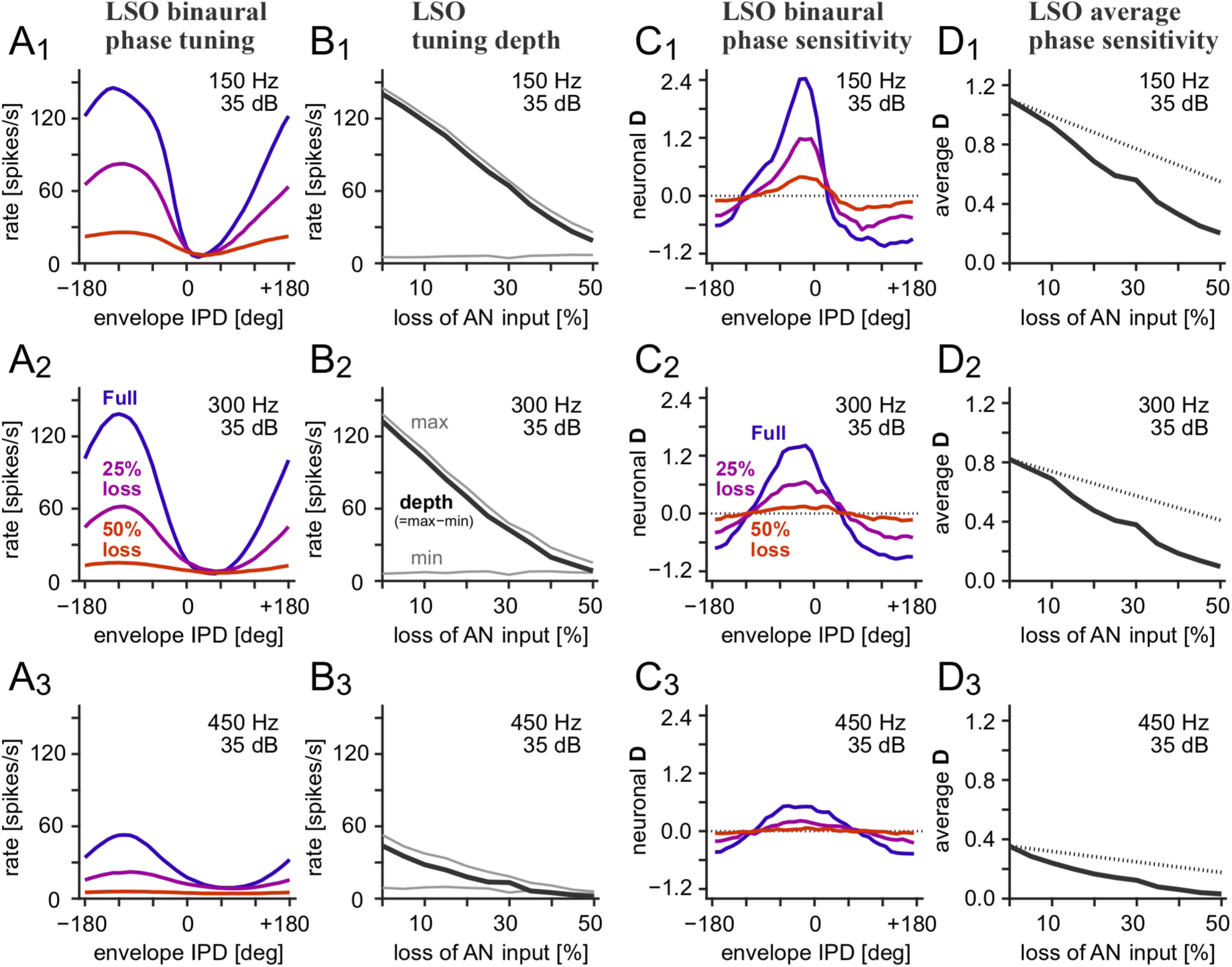
| Binaural phase tuning of LSO across degrees of input loss. Responses were simulated at 35 dB SPL with matched fractional AN input loss in both bushy-cell pathways and no additional jitter. Rows 1–3 show envelope frequencies of 150, 300 and 450 Hz. **(A_1_–A_3_)** LSO rate versus envelope IPD. Blue, purple, and orange curves indicate intact input, 25% loss and 50% loss, respectively. A positive and negative IPD indicates the contralateral sound leading and lagging ipsilateral sound, respectively. **(B_1_–B_3_)** Maximum and minimum LSO rates across the phase-tuning curve (thin gray) and tuning depth (defined as maximum minus minimum rate: thick black) versus input loss. **(C_1_–C_3_)** LSO neuronal discriminability **D** versus envelope IPD calculated for the tuning curves in A_1_–A_3_. **(D_1_–D_3_)** Root-mean-square average of LSO neuronal **D** calculated over the envelope IPD tested (between –180 and +180 degrees). Dotted lines indicate a decay proportional to AN input loss.

To assess the separation of responses to small changes in envelope IPDs, we calculated neural discriminability **D**, defined as the difference in mean LSO firing rates at IPDs separated by 10 degrees divided by the average of their response standard deviations. Its sign indicates the direction of the local rate change, and its magnitude reflects response separation relative to trial-to-trial variability. With intact input, pronounced positive and negative excursions occurred along opposing flanks of the phase-tuning curves (Fig. 4C1-C3). Removing 25% of AN inputs reduced these excursions, and 50% loss left little residual IPD discriminability at 300 and 450 Hz, with greater preservation at 150 Hz. Root-mean-square-averaged **D** across the tested IPDs declined more steeply than the reference proportional to surviving AN inputs, with larger fractional reductions at higher modulation frequencies (Fig. 4D1-D3). Thus, largely preserved phase locking of bushy cells coexisted with weaker neural discriminability of IPD in LSO.

### Jitter in bushy cell spike timing degrades phase locking without removing spikes

To isolate timing disruption, we retained all AN inputs and added jitter to bushy cell output spikes (Fig. 5). Rate-level functions with no added jitter, 0.4-ms jitter and 0.8-ms jitter closely overlapped in both populations at all three envelope frequencies (Fig. 5A1-A3,B1-B3), as expected when spike times were displaced without deleting spikes.

**Fig. 5.**
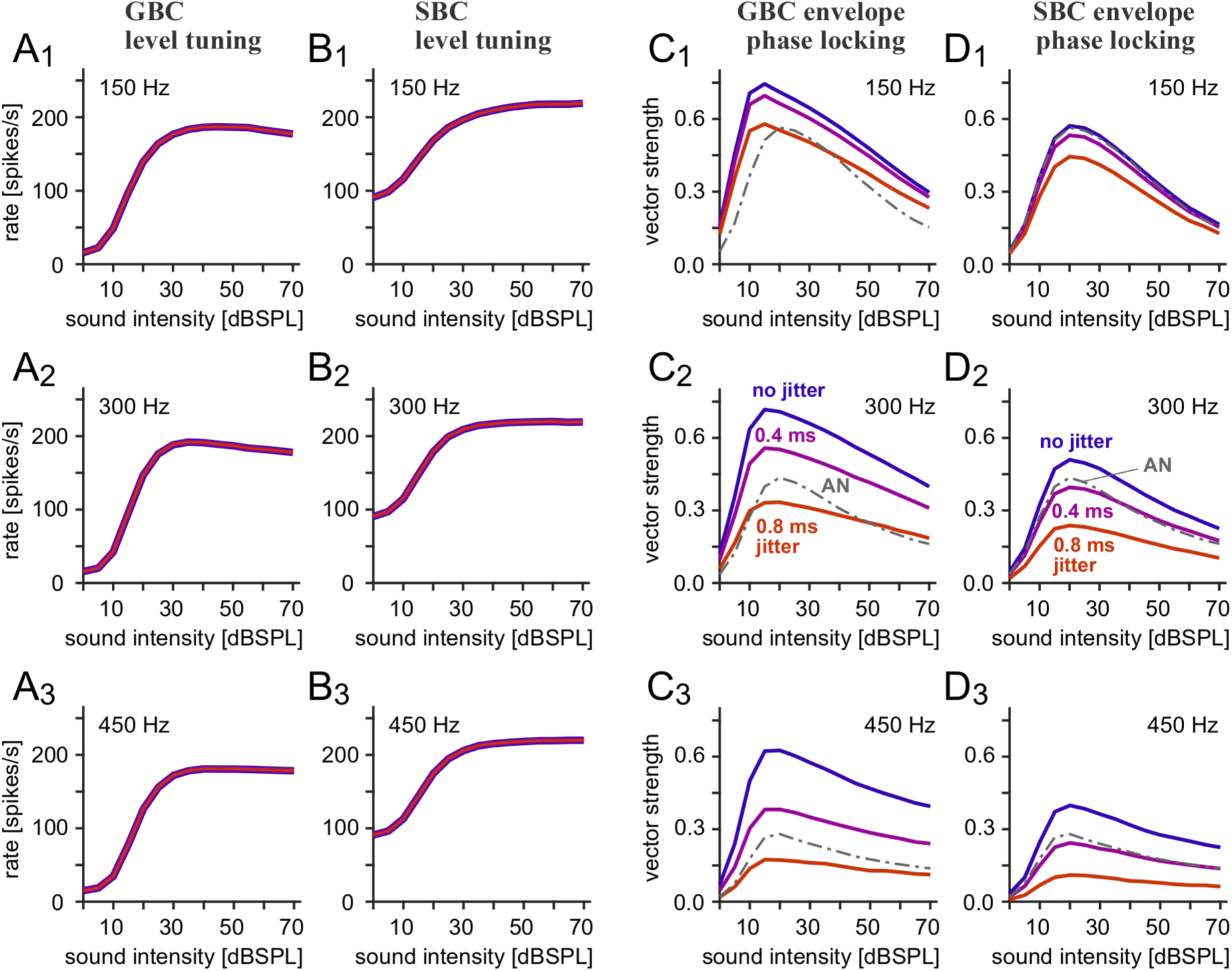
| Bushy-cell firing rates and phase locking with added output jitter. With all AN inputs retained, responses are plotted against sound level, with rows 1–3 showing envelope frequencies of 150, 300 and 450 Hz. **(A_1_–A_3_)** GBC rates. **(B_1_–B_3_)** SBC rates. **(C_1_–C_3_)** GBC output VS. **(D_1_–D_3_)** SBC output VS. Blue, purple, and orange indicate no added jitter, 0.4-ms jitter and 0.8-ms jitter, respectively. Gray dash-dotted curves show the unperturbed AN phase locking reference where visible. Jitter was added to bushy-cell output times before delivery to the LSO; VS therefore describes temporally perturbed outputs rather than an altered spike-generation mechanism. Rate curves closely overlap.

VS decreased with increasing jitter, with progressively larger effects at higher envelope frequencies (Fig. 5C1-C3,D1-D3). At 150 Hz, 0.4-ms jitter caused a small decrease, and substantial phase locking remained with 0.8-ms jitter. The same perturbations reduced VS more severely at 300 and 450 Hz, where a fixed jitter occupies a larger fraction of the envelope cycle. With 0.8-ms jitter, bushy cell VS could fall below the unperturbed AN reference at these frequencies. The characteristic level-dependent rise and fall in VS remained visible but was attenuated. Temporal dispersion thus eroded the temporal precision initially gained through bushy cell processing while preserving overall spike count.

### Faster envelope fluctuations increase vulnerability to timing dispersion

We varied temporal jitter over a wider range at 35 dB SPL to determine its consequences for binaural coding. Bushy cell firing rates remained constant (Fig. 6A1-A3), whereas VS decreased in both populations (Fig. 6B1-B3). The decrease was gradual at 150 Hz and occurred over progressively smaller jitter values at 300 and 450 Hz. GBCs retained stronger phase locking than SBCs, with the same frequency ordering in both populations.

**Fig. 6.**
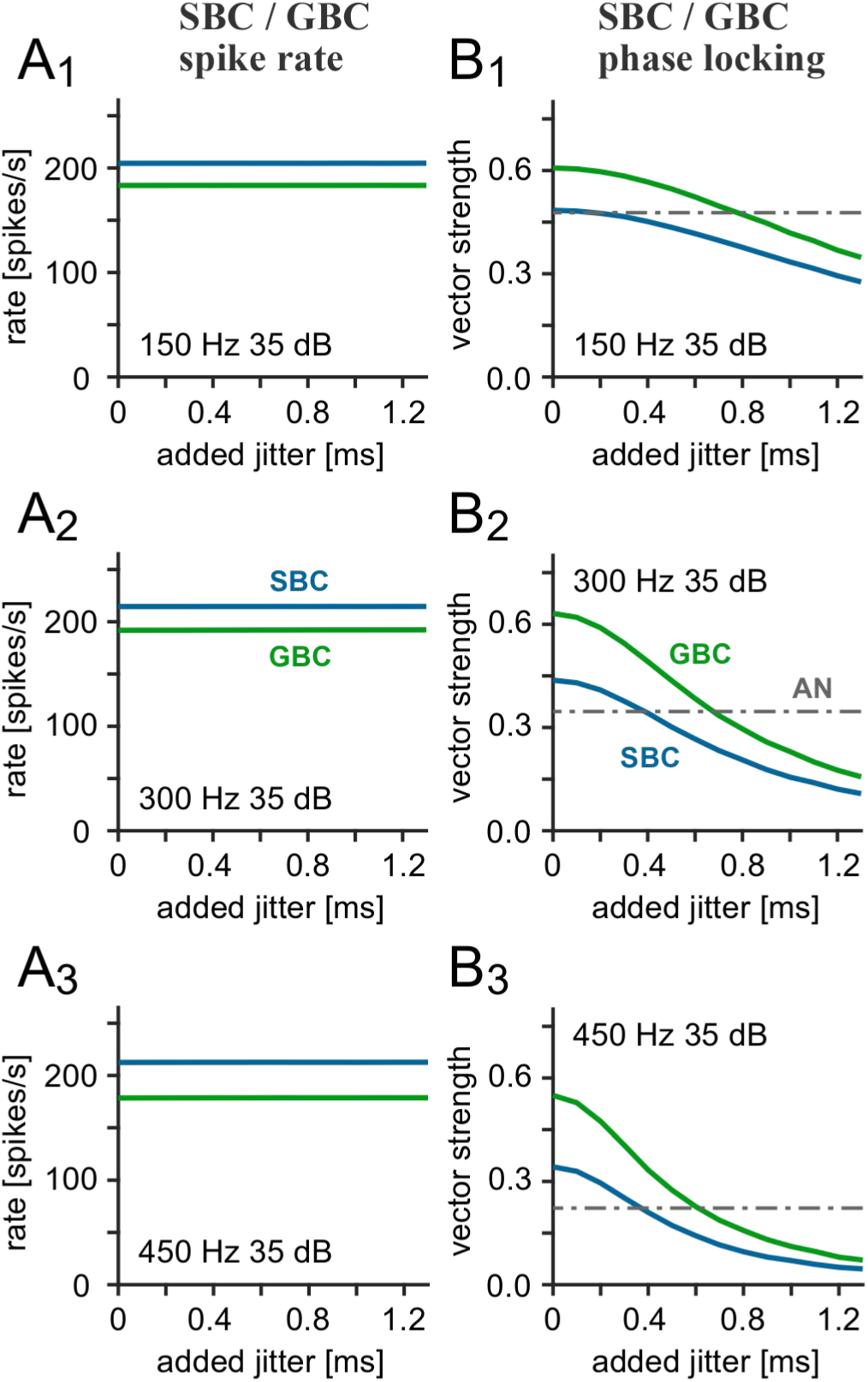
| Bushy-cell responses across jitter magnitudes. Temporal jitter was added to spiking output of bushy cells, while all AN inputs were retained intact. Responses were simulated at 35 dB SPL. Rows 1–3 show envelope frequencies of 150, 300 and 450 Hz, respectively. **(A_1_–A_3_)** Bushy-cell rates versus added jitter; light blue and light green indicate SBCs and GBCs, respectively. **(B_1_–B_3_)** Corresponding output VS. Gray dash-dotted lines indicate the unperturbed AN reference, which was not subjected to output jitter.

LSO tuning to IPD followed this frequency dependence (Fig. 7A1-A3). At 150 Hz, clear phase-dependent firing remained with 0.4– and 0.8-ms jitter, although the peak declined. At 300 Hz, 0.4-ms jitter substantially attenuated the peak and 0.8-ms jitter left a small phase-dependent response. At 450 Hz, tuning was markedly shallower with 0.4-ms jitter and nearly flat with 0.8– ms jitter. Response maxima and minima converged as jitter increased, predominantly through a falling maximum (Fig. 7B1-B3). Tuning depth was relatively preserved up to approximately 0.2 ms, then approached zero as jitter increased. Higher envelope frequencies showed this decline at smaller jitter values.

**Fig. 7.**
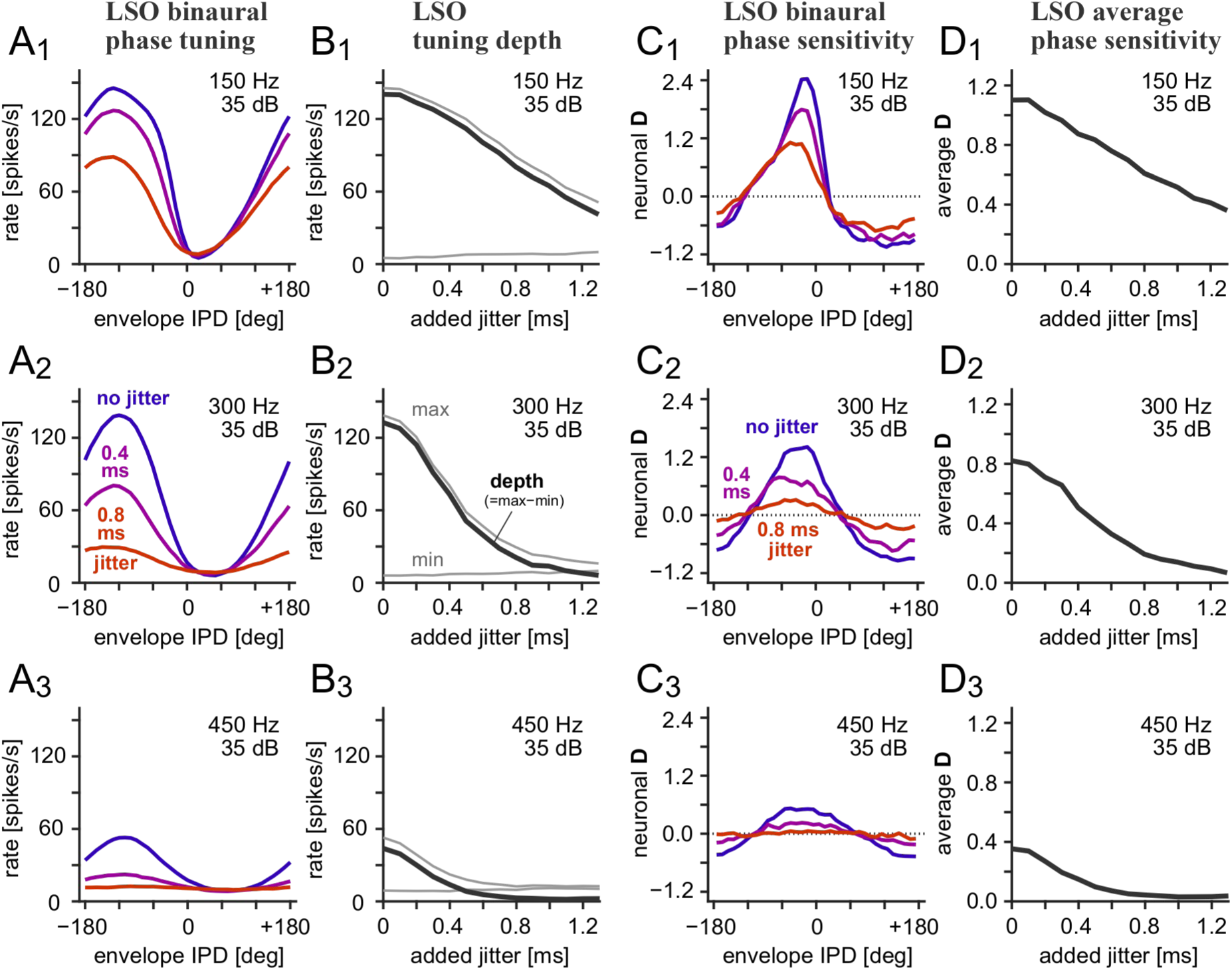
| Binaural phase tuning of LSO across jitter magnitudes. All AN inputs were retained and responses were simulated at 35 dB SPL. Temporal jitter was added to spiking output of bushy cells. Rows 1–3 show envelope frequencies of 150, 300 and 450 Hz, respectively. **(A_1_–A_3_)** LSO phase tuning with no added jitter (blue), 0.4-ms jitter (purple) and 0.8-ms jitter (orange). The same jitter condition was applied to both pathways. A positive IPD indicates the contralateral sound leading ipsilateral sound**. (B_1_–B_3_)** Maximum and minimum LSO rates (thin gray) and tuning depth (maximum minus minimum: thick black) versus jitter. **(C_1_–C_3_)** LSO neuronal discriminability **D** versus envelope IPD calculated for the tuning curves in A_1_–A_3_. **(D_1_–D_3_)** Root-mean-square average of LSO neuronal **D** calculated over the envelope IPD tested (between –180 and +180 degrees).

Jitter also brought the prominent positive and negative excursions of discriminability **D** closer to zero (Fig. 7C1-C3). Substantial discriminability remained at 150 Hz with both perturbations, whereas reductions were larger at 300 Hz and little residual discriminability remained at 450 Hz with 0.8-ms jitter. Average D was relatively preserved at the smallest jitter values and declined with increasing dispersion, with larger fractional reductions at higher frequencies (Fig. 7D1-D3). These reductions in neighboring-response separation occurred while bushy cell firing rates were preserved.

### Combined input loss and spike timing dispersion further weaken binaural coding

We examined the joint effects of deafferentation and timing disruption by combining 25% AN input loss with 0.4-ms output jitter. This further attenuated IPD tuning in LSO relative to either perturbation alone (Fig. 8A1-A3). At 150 Hz, jitter alone had a comparatively small effect and input loss accounted for most attenuation. At 300 Hz, both perturbations substantially reduced phase-dependent firing. At 450 Hz, either perturbation left little IPD modulation and their combination further weakened it.

**Fig. 8.**
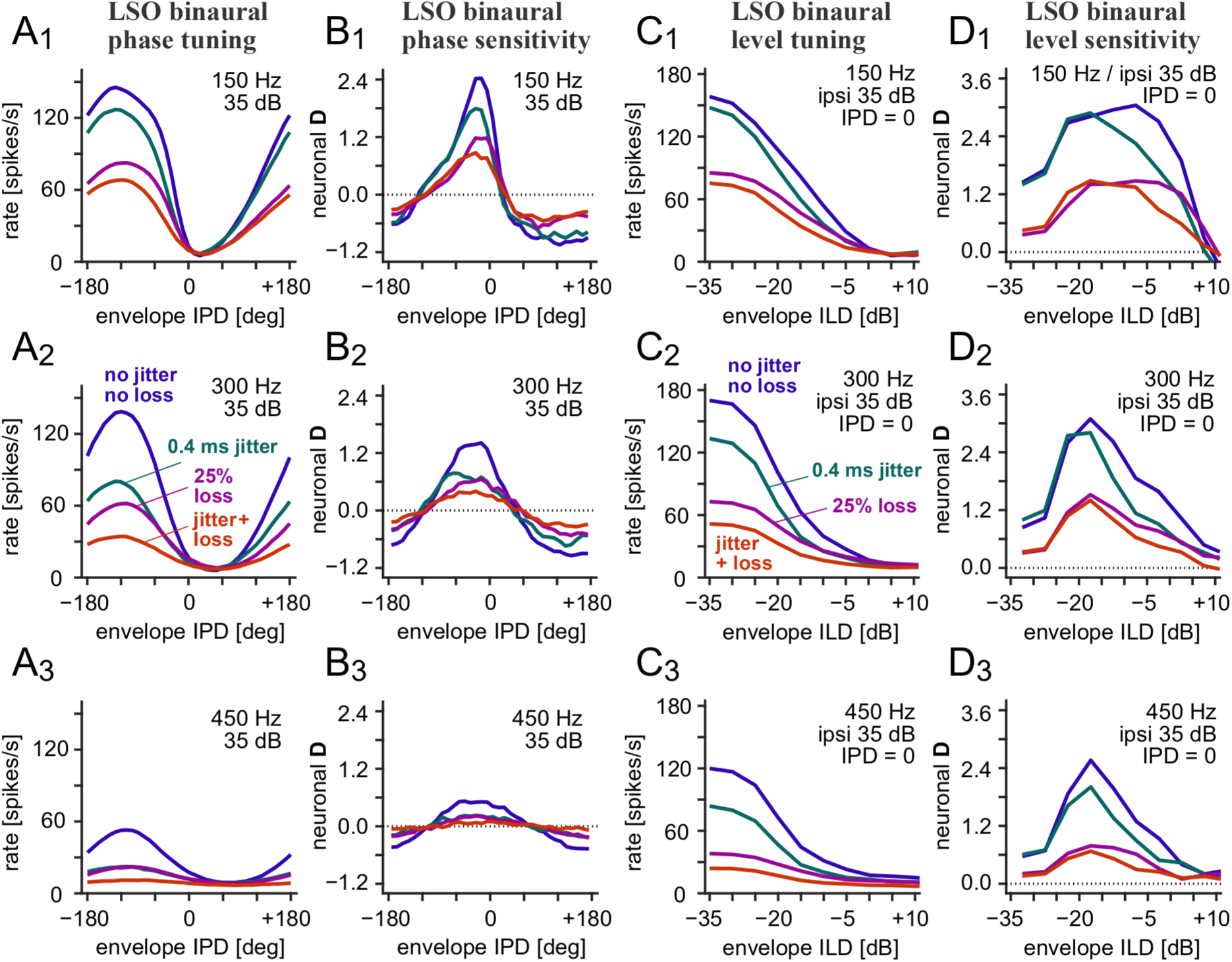
| Binaural phase and level tuning with separate and combined perturbations. Rows 1–3 show envelope frequencies of 150, 300 and 450 Hz, respectively. Simulated conditions are: intact AN inputs without added jitter (blue), 25% AN input loss without jitter (purple), 0.4-ms output jitter with intact AN inputs (teal), and combined input loss and jitter (orange). Each perturbation was applied simultaneously to both bushy-cell pathways. **(A_1_–A_3_)** LSO rate versus envelope IPD with both ears at 35 dB SPL. A positive IPD indicates the contralateral sound leading ipsilateral sound. **(B_1_–B_3_)** LSO neuronal discriminability **D** versus envelope IPD calculated for the tuning curves in A_1_–A_3_. **(C_1_–C_3_)** LSO rate versus ILD for AM sounds with ipsilateral level fixed at 35 dB SPL and envelope IPD at zero. Contralateral level was varied to produce the plotted ILDs. A positive and negative ILD indicates the contralateral sound being stronger and weaker than ipsilateral sound, respectively. These level-tuning functions concern AM sounds, not unmodulated tones. **(D_1_–D_3_)** LSO neuronal discriminability **D** versus ILD calculated for the tuning curves in C_1_–C_3_.

At 300 Hz, tuning depth fell from 133 to 54 spikes/s with input loss and to 73 spikes/s with jitter, retaining approximately 41% and 55% of intact depth. Their combination retained approximately 20% (27 spikes/s) of intact depth, close to the product of the individual retained fractions (23%). A similar relation was apparent at 450 Hz. These comparisons suggest approximately multiplicative attenuation of IPD tuning depth at this operating point.

The combined perturbation also produced the greatest reduction in the prominent extremes of IPD neural discriminability (Fig. 8B1-B3). At 150 Hz, input loss accounted for most of the reduction in **D** magnitude, with additional attenuation by imposed jitter. At 300 Hz, both input loss and jitter reduced the positive and negative excursions, and their combination compressed the discriminability profile further toward zero. At 450 Hz, each perturbation alone left limited phase discriminability and the combined condition produced the weakest residual profile.

Both perturbations compressed sound level tuning for AM sounds (Fig. 8C1-C3). With the ipsilateral level fixed at 35 dB SPL and envelope IPD at zero, intact LSO firing decreased as ILD increased. Input loss and jitter each reduced the high rate part of the tuning curve, and their combination further decreased the response range. We calculated local ILD discriminability between responses separated by 5 dB (Fig. 8D1-D3). Jitter alone preserved substantial discriminability, although its effect varied across ILDs. At 150 Hz, the region of greatest sensitivity shifted toward more negative ILDs, where **D** remained comparable to intact. Input loss reduced peak ILD discriminability more strongly, and the combined perturbation attenuated **D** over much of the tested range. At 450 Hz, jitter retained a pronounced ILD-discriminability peak despite strongly reduced IPD discriminability, whereas input loss and the combined perturbation produced much smaller values of ILD discriminability. Temporal dispersion therefore had different consequences depending on the binaural cue, IPD or ILD.

### Pathway-selective input loss dissociates firing rate from binaural interaction

Finally, we applied 50% AN input loss to GBCs, SBCs or both populations without adding jitter (Fig. 9). GBC-selective loss reduced inhibitory drive while SBC excitation remained intact, increasing LSO firing and elevating the trough of phase tuning (Fig. 9A1-A3). At 300 Hz, rates exceeded approximately 100 spikes/s even at IPDs that nearly silenced the intact model. LSO rates also remained elevated at ILDs that strongly suppressed intact responses (Fig. 9C1-C3). At 450 Hz, substantial activity persisted throughout the tested ranges despite weak binaural modulation. SBC-selective loss produced the opposite rate change, reducing excitation while GBC-driven inhibition remained intact. Small residual responses remained at 150 Hz, with especially weak responses at 300 and 450 Hz. Simultaneous loss produced low-rate, weakly tuned responses. However, the accompanying reduction in inhibition partly offset the rate decrease caused by SBC loss, so combined-loss rates did not necessarily fall below SBC-only rates.

**Fig. 9.**
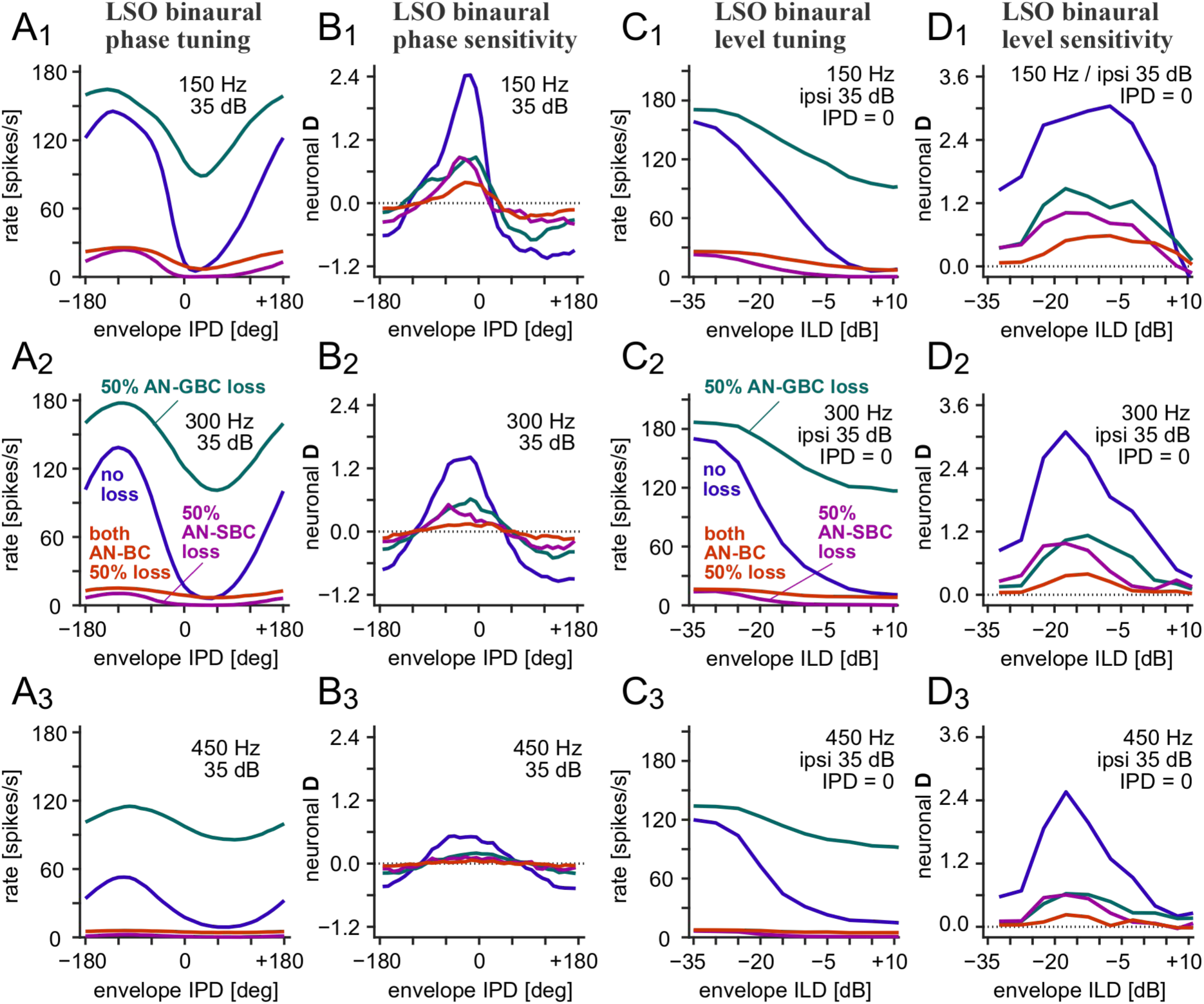
| Binaural phase and level tuning after pathway-selective input loss. Rows 1–3 show envelope frequencies of 150, 300 and 450 Hz, with no output jitter added. Blue indicates intact AN input. Teal indicates 50% AN input loss restricted to GBCs, reducing inhibitory drive while preserving SBC excitation. Purple indicates 50% AN loss restricted to SBCs, reducing excitation while preserving the GBC pathway. Orange indicates 50% loss in both populations (“both AN-BC”). **(A_1_–A_3_)** LSO rate versus envelope IPD with both ears at 35 dB SPL. A positive IPD indicates the contralateral sound leading ipsilateral sound. **(B_1_–B_3_)** LSO neuronal discriminability **D** versus envelope IPD calculated for the tuning curves in A_1_–A_3_. **(C_1_–C_3_)** LSO rate versus ILD for AM sounds with ipsilateral level at 35 dB SPL and envelope IPD at zero. A positive ILD indicates the contralateral sound being stronger than ipsilateral sound. The manipulation removes functional AN inputs to bushy cells, without reducing the SBC-to-LSO or GBC/MNTB-to-LSO contacts. **(D_1_–D_3_)** LSO neuronal discriminability **D** versus ILD calculated for the tuning curves in C_1_–C_3_.

Pathway-selective loss reduced neural discriminability despite these opposing changes in overall firing (Fig. 9B1-B3,D1-D3). GBC-selective loss (teal) reduced prominent IPD-discriminability extrema and peak ILD discriminability. SBC-selective loss (purple) also weakened discriminability, although appreciable local sensitivity remained at some disparities, particularly at 150 Hz. The two selective-manipulation profiles overlapped in some regions despite their markedly different firing rates. Simultaneous loss produced less prominent **D** values than either selective manipulation, even though reduced inhibition partly offset the rate decrease caused by SBC loss. At 450 Hz, both selective manipulations retained modest ILD discriminability, whereas simultaneous loss left only weak residual values. Thus, reduced local binaural discrimination accompanied either increased or decreased LSO activity.

## Discussion

The central result of the present study is a dissociation between precise monaural spike timing and effective binaural computation. Removing functional AN inputs to simulate cochlear synaptopathy left bushy cell phase locking largely intact, yet disproportionately reduced LSO binaural tuning (Figs. 2-4). Timing jitter weakened the same computation without removing spikes (Figs. 5 and 7), and combined perturbations further compressed the response range (Fig. 8). Reduced neural discriminability demonstrates that preserved bushy cell phase locking can coexist with less separable LSO responses to small changes in envelope phases after accounting for trial-to-trial variability (Fig. 4). Temporal dispersion further reduced discriminability (Figs. 7 and 8), linking spike availability and timing to the quality of binaural representations. The temporal sharpening provided by bushy cells therefore cannot fully protect downstream computation when too few spikes remain. Effective binaural interaction requires sufficient spikes arriving in combinations that excite or inhibit the LSO.

### Input availability limits coincidence sensitive processing

Peripheral input loss cannot be propagated through the brainstem simply as a uniform scaling factor. GBCs and SBCs receive the same total input count in the model but differ in synaptic weighting and threshold-dependent integration [8, 9], producing unequal fractional losses of their outputs. At the LSO, excitation must reach threshold within a short window while interacting with temporally structured inhibition [11, 13]. Fewer inputs reduce the probability of effective coincidences even when preferred firing phases are preserved. The coincidence-sensitive transformations at the bushy cells that improve temporal representation in the intact pathway [7, 8] can therefore make downstream computation vulnerable to depleted input, leading to more intense functional consequences of lost convergence than anatomical deafferentation.

Comparisons with prior synaptopathy models depend on the response measures and afferent assumptions. Budak and colleagues demonstrated impaired ITD tuning in MSO after peripheral synaptopathy or myelin defects [21]. Their relative vector strength equals conventional VS multiplied by the test-to-control spike count ratio, so it can decline without poorer phase locking of surviving spikes. The preserved conventional VS we show here is therefore compatible with their findings. Their greater SBC than GBC susceptibility contrasts with the present spike rate ordering, identifying afferent composition and integration assumptions as determinants of vulnerability rather than supporting a universal hierarchy between bushy cell types.

Physiological findings reinforce the importance of matched stimulus conditions. Hockley and colleagues found reduced suprathreshold tone-in-noise responses in cochlear nucleus small cells and primary-like units after confirmed synaptopathy, but preserved mean rate level functions for tones in quiet [25]. Cochlear synapse loss therefore need not produce the stimulus rate decrease obtained here. Stimulus statistics, fiber selectivity and compensation can determine whether reduced peripheral drive is expressed in central output. Our simulations isolate one transformation of effective input loss whose quantitative correspondence to physiological synaptopathy requires empirical testing.

### Timing dispersion and pathway imbalance provide distinct routes to binaural coding deficits

A different failure mode emerges when available spikes are dispersed in time. At higher envelope frequencies, a fixed jitter occupies a larger fraction of the stimulus cycle and broadens the distribution of spike phases. Greater vulnerability to jitter for faster envelopes therefore concerns temporal demands rather than necessarily carrier frequency or cochlear place. Jitter was imposed after spike generation, making rate preservation a controlled feature of the model. An in vitro study in a spontaneous dysmyelinating rat phenotype reported approximately 0.15-ms conduction jitter between afferent fibers and MNTB neurons [16]. Conductance-clamp recordings and modeling also indicate natural variation of approximately 0.1–0.2 ms in bushy cell spike generation [26]. These observations establish plausible sources of timing variability, although the imposed jitter magnitudes require direct physiological calibration. Axonal changes can also alter mean delays between pathways, shifting binaural tuning, or produce heterogeneous delays among axons that disperse coincidences [15, 22]. Separating common delays among inputs, pathway offsets, and dispersion will be needed to determine how specific myelin abnormalities eventually affect binaural computation. Our simulations here isolate one transmission defect within this broader range of abnormalities.

Pathway selective loss shows why the sign of a central rate change is insufficient to identify preserved function or the site of injury. Removing AN input to GBCs reduced inhibitory drive and elevated LSO firing, whereas removing input to SBCs reduced excitation and lowered firing. Both reduced prominent IPD neural discriminability magnitudes (Fig. 9). Peripheral deafferentation can thus produce central disinhibition without a frank loss of inhibitory neurons. Earlier LSO models showed resilience to direct inhibitory input loss when surviving synapses were strengthened [23], and central gain can restore aspects of rate-based responses while temporal deficits persist [27]. The results of neural discriminability measure **D** extend this distinction. Simultaneous loss partly offset the rate decrease caused by SBC loss while producing smaller prominent **D** values than either selective manipulation. Functional compensation should therefore be evaluated through separation of responses to task-relevant cue differences relative to variability, including the cue range supporting sensitivity.

Combined input loss and jitter produced approximately multiplicative attenuation of tuning depth at the tested operating point (Fig. 8). Neural **D** further revealed cue-dependent consequences. Relative to the corresponding intact profiles, 0.4-ms jitter strongly reduced envelope-phase discriminability at 450 Hz while preserving a pronounced region of ILD discriminability (Fig. 8B3,D3). The consequences therefore depend on the binaural cue, IPD or ILD, and their operating ranges. Since AM level responses are assembled from temporally structured inputs, timing abnormalities can weaken representations of phase and level cues to different degrees.

### Reduced bushy cell activity provides a candidate link to central myelin dysfunction

The two functional impairments could share a biological origin. AN input loss reduces bushy cell output and the activity propagated along bushy cell axons, particularly in the modeled GBC pathway (Fig. 2A1-A3). If sustained during everyday stimulation, this reduction could weaken activity-dependent signals regulating myelin. Preserved phase locking would not prevent this effect, because fewer action potentials can retain precise timing. The model thus identifies a neuronal intermediate through which peripheral deafferentation might influence central spike transmission, while leaving open whether the activity decrease is sufficiently persistent or large to alter myelin in vivo.

Auditory deprivation studies provide anatomical and functional precedents. Conductive hearing loss in adult mice reduced myelin thickness in the largest trapezoid body fibers [17]. Active and deprived GBC axons within the same bundles differ in myelination during postnatal auditory development, indicating regulation at the level of individual axons [18]. Following eight weeks of developmental unilateral earplugging in guinea pigs, binaural brainstem responses and behavioral spatial acuity remain impaired after earplug removal despite recovered hearing thresholds [38]. These results establish adult sensitivity to reduced auditory drive, developmental axon specificity, and persistent functional consequences of deprivation. None of these studies used selective cochlear synaptopathy, and the developmental myelination result should not be interpreted as destruction of established adult myelin. Together, however, they support altered myelination of bushy cell axons as a candidate contributor to deprivation-associated binaural dysfunction.

Applied to cochlear synaptopathy, this comparison motivates a shared downstream mechanism in which reduced bushy cell output alters the activity available to regulate central myelin. Establishing that mechanism, however, requires evidence connecting peripheral synapse loss to persistent changes in axonal structure and transmission, while distinguishing transmission abnormalities from altered synaptic integration. This distinction is essential because both conductive loss and cochlear synaptopathy reduce auditory drive in the brainstem yet through different mechanisms.

Several cellular mechanisms could mediate this link. Neuronal stimulation promotes oligodendrocyte generation and axonal myelination, whereas developmental suppression of selected axons reduces their probability of myelination [19, 20]. In the auditory brainstem, some premyelinating oligodendrocytes receive glutamatergic input and generate sodium-channel-dependent spikes, with reduced excitability impairing axon–glia interactions and myelination [28]. Neuronal input also recruits calcium signals in developing oligodendrocytes [29]. Culture experiments link vesicular release at nonsynaptic axon–glia contacts to calcium signaling, myelin protein synthesis and preferential myelination of active axons [30]. These findings identify possible neural activity sensors, but leave unresolved whether adult bushy cell output is monitored through mean firing rate, burst structure or another activity feature.

Activity-dependent explanations must distinguish failed formation or repair from deterioration of an established myelin sheath. Adult cortical oligodendrocytes that successfully integrate are highly stable, and sensory enrichment primarily promotes new oligodendrocyte integration rather than changing existing sheath length [31]. Reduced activity therefore cannot simply be equated with mature oligodendrocyte death. Blocking neuronal activity or glutamatergic signaling, however, impairs oligodendrocyte differentiation and remyelination after an experimental demyelinating lesion [32]. Persistently reduced bushy cell activity might thus favor tract-specific thinning or compromise repair when myelin is independently injured. Synaptopathy could consequently prolong a central myelin deficit without being its sole initiating cause.

We propose a conditional progression from cochlear synapse loss through reduced bushy cell output to inadequate myelin regulation and less reliable temporal transmission of surviving spikes. Myelin thinning, altered nodal organization or incomplete repair could contribute. In aged gerbils, abnormal MNTB-afferent myelination and oligodendrocyte loss was shown to accompany altered binaural brainstem responses and poorer spatial hearing [33], supporting the relevance of this pathway. The present simulations contain neither glial dynamics nor a model linking firing rate to sheath structure, but demonstrate the functional cost of reduced drive combined with temporal dispersion. If the proposed transition occurs, binaural coding could continue to deteriorate even after peripheral synapse loss stabilizes. Maintaining bushy cell activity or myelin could prevent that additional impairment, although noise exposure and aging could also damage synapses and myelin through partly independent processes.

### Restoring transmission timing alone does not resolve the consequences of synaptopathy

Separating reduced afferent drive from impaired transmission timing has important therapeutic implications. Correcting myelin-related timing abnormalities alone would not be expected to normalize binaural coding when cochlear deafferentation persists. At 300-Hz modulation, combined 25% AN input loss and 0.4-ms jitter retained approximately 20% of intact tuning depth, whereas input loss alone retained approximately 40% (Fig. 8). An idealized intervention eliminating added jitter while holding afferent loss and other circuit properties fixed would therefore improve responses but leave tuning depth approximately 60% below intact. Neural discriminability extends this prediction. Prominent IPD-discriminability magnitudes and peak ILD discriminability remained substantially below intact values in the input-loss-only condition (Fig. 8B2,D2). Restoring spike time precision alone cannot replace the missing spikes required for effective coincidence-sensitive processing by the LSO.

These findings support cochlear deafferentation and central transmission dysfunction as complementary therapeutic targets. Where both coexist, improving myelination may alleviate timing-related impairment while leaving a substantial binaural deficit from reduced input. Conversely, restoring afferent inputs could leave residual impairment if transmission timing remains abnormal. Degraded binaural representations could limit spatial discrimination and the benefit of spatial separation for understanding speech among competing sources despite normal or near-normal thresholds. Yet these behavioral consequences remain predictions. Our model motivates interventions that restore or functionally compensate for afferent drive lost to synaptopathy, together with preservation or repair of central transmission where needed. Their effectiveness in vivo will depend on the accompanying biological lesions and compensatory responses.

### Implications for physiological biomarkers and spatial hearing

Reduced periodic population activity can reflect fewer spikes, poorer phase locking or both. Associations between synaptopathy and temporal envelope response deficits [34] should be interpreted in that light, while recognizing that scalp-recorded envelope-following responses also depend on generator composition and summation. The binaural interaction component (BIC) of the auditory brainstem response provides a complementary circuit measure, because physiological comparisons and modeling implicate the LSO in its generation [35, 36]. Insufficient or mistimed inputs could both reduce binaural interaction, so an abnormal BIC would not uniquely identify damaged inhibitory synapses, but could still serve as a proxy for the impaired temporal precision of monaural inputs.

Neither evoked potential waveforms nor behavioral performance were simulated here. Relating the sustained AM responses modeled here to click-evoked brainstem potentials requires an explicit forward model [37]. Combining population responses with identified proximal and distal bushy cell recordings could distinguish reduced spike generation from an additional transmission deficit. Such measurements would help connect the separate manipulations in this model to the lesions underlying altered physiological markers.

The neural **D** discrimination metric incorporates local mean spike rate differences and trial-to-trial response variability [23], providing a closer connection between circuit responses and binaural behavior. Reduced **D** magnitudes indicate weaker neural ability to separate small changes in binaural cue values (Figs. 4, 7-9). Behavioral thresholds additionally require evaluation across cue increments, an explicit decision rule [39] and specification of downstream population pooling, which can preserve sensitivity despite substantial neural loss [40]. A population readout incorporating correlations and naturally occurring disparities would help quantify spatial acuity predictions, while speech understanding predictions require task-appropriate stimuli.

Monaural and binaural tasks need not be affected equally. Residual phase locking could support some modulation or periodicity judgments, whereas binaural comparisons additionally require appropriate timing and balance across converging pathways. This predicted dissociation should be tested with matched stimuli. Human associations between subcortical temporal coding and difficult listening [1, 41] are compatible with the mechanism, while absent electrophysiological evidence for synaptopathy in other listeners [5] cautions against a single-cause account. Middle-aged gerbils can also show impaired binaural masking level differences without reduced apical synapse counts [42]. Generalization to low-frequency fine-structure tasks, age-related deficits or speech understanding therefore requires further evidence.

### Model limitations and experimental tests

The simulated AN fibers all have high spontaneous rates, whereas low-spontaneous-rate fibers show selective vulnerability in a guinea pig synaptopathy model [43]. Histological synapse loss therefore cannot be mapped directly onto active model inputs removed. The manipulation represents withdrawal of functional afferent drive rather than anatomical elimination of cochlear-nucleus endbulbs or boutons. Robustness remains to be established across synaptic-strength heterogeneity, afferent correlations, homeostatic plasticity, transmission failures, realistic afferent mixtures and lesion patterns. Background noise and transient stimuli are also important because abrupt acoustic changes recruit mechanisms beyond the sustained responses represented here [44].

A decisive test would quantify cochlear innervation, identified bushy cell firing, axonal myelin and binaural behavioral function longitudinally, then manipulate bushy cell activity while keeping peripheral synapse loss approximately constant. Preservation of distal timing and myelin with maintained activity would support the proposed link, whereas continued deterioration would favor additional mechanisms. Electron microscopy, nodal measurements and oligodendrocyte-lineage analyses could distinguish thinning, impaired maturation and failed repair. Separate SBC and GBC measurements would test pathway specificity, and established gerbil spatial hearing assays [45] could connect these changes to behavior. Comparing recovery of drive, transmission timing and both together would test whether correcting one component leaves residual impairment attributable to the other.

In summary, our simulations demonstrate that preserved monaural spike precision does not guarantee effective binaural computation. Coincidence-sensitive processing can transform partial deafferentation into disproportionately weak LSO responses and neural discrimination of binaural cues, with temporal dispersion compounding the impairment. The model separates insufficient spike supply from impaired spike timing transmission while motivating a testable connection through activity-dependent myelin regulation. Longitudinal experiments will be necessary to establish whether cochlear synaptic injury acquires secondary central consequences. Restoring or compensating for lost afferent drive remains a treatment objective even when accompanying timing dysfunction can be corrected.

## Methods

### AN-BC-LSO model

The simulation framework in this study here was adopted from a recent modeling paper [8], in which detailed descriptions on the structure of the model and selection of the parameters are provided. Here we briefly review the model and describe relevant modifications. Spiking trains of high-spontaneous-rate AN fibers (70 spikes/s) simulated with an auditory periphery model [24] were fed into contralaterally driven GBC model and ispsilaterally driven SBC model [8, 9]. The GBC model receives 20 subthreshold synaptic inputs of equal size, while the SBC model receive two suprathreshold inputs and 18 subthreshold inputs, respectively mimicking endbulb and bouton inputs (Fig. 1B). The LSO model receives 20 excitatory inputs from SBCs and 8 inhibitory inputs from GBCs, assuming that MNTB neurons that are located between GBCs and LSO is a faithful sign inverter. The numbers of inputs to LSO were adjusted from those used previously [8], for the sake of easier implementation of systematic input reduction described below; the simulated binaural response characteristics of the LSO model remain unaffected with this modification.

Both bushy cell models operate as coincidence detectors of synaptic inputs (Fig. 1C,D). A sufficient number of AN spikes arriving within a short time window lead to a generation of an output spike, while poorly synchronized inputs are rejected by the adaptive threshold. Membrane parameters were identical from the previous work [8]. The LSO model operates in a similar way, but with inhibitory inputs and without threshold adaptation (Fig. 1E). Each inhibitory input cancels two excitatory inputs arriving within the inhibitory time window; this excitatory-inhibitory interaction is the basis for binaural time– and level-difference tuning [13].

### Reduction of AN inputs to bushy cells

Cochlear synaptopathy, or the loss of AN fibers projecting to neurons in the cochlear nuclei, was modeled as a reduction of AN inputs to bushy cells. For the GBC model, in which all inputs have the same size, the number of AN projections per cell was simply reduced from 20 (intact condition) down to ten (50% loss) with a step of one (corresponding to 5%). For the SBC model, in which endbulb and bouton synapses coexist, the modification of input numbers was determined in a way that the total number of each synapse type eventually feeding the target LSO cell is reduced proportionally to the loss rate (Table 1). In the case of 50% loss, for example, the resulting numbers are one endbulb input and nine bouton inputs, both of which are a half of the original values. For a loss rate of 25%, a half of SBCs projecting to LSO have one endbulb (and 14 boutons), while the other SBCs have two endbulbs (and 13 boutons); both leading to 15 synaptic inputs in total (i.e., 75% of the original count).

**Table 1.** Number of SBC inputs to LSO with loss of AN fibers. The input configuration is designed such that the average number of endbulb AN inputs per SBC is always 10% of the total number of AN inputs per SBC. The total number of SBC inputs per LSO is fixed to 20, regardless of AN loss.

| **AN loss rate** | **Input configuration to LSO** | **Total number of AN inputs per SBC** |
|---|---|---|
| **0% (intact)** | 20 SBCs with two endbulbs and 18 boutons | **20** |
| 5% | 2 SBCs with one endbulb and 18 boutons +<br>18 SBCs with two endbulbs and 17 boutons | 19 |
| 10% | 4 SBCs with one endbulb and 17 boutons +<br>16 SBCs with two endbulbs and 16 boutons | 18 |
| 15% | 6 SBCs with one endbulb and 16 boutons +<br>14 SBCs with two endbulbs and 15 boutons | 17 |
| 20% | 8 SBCs with one endbulb and 15 boutons +<br>12 SBCs with two endbulbs and 14 boutons | 16 |
| **25%** | 10 SBCs with one endbulb and 14 boutons +<br>10 SBCs with two endbulbs and 13 boutons | **15** |
| 30% | 12 SBCs with one endbulb and 13 boutons +<br>8 SBCs with two endbulbs and 12 boutons | 14 |
| 35% | 14 SBCs with one endbulb and 12 boutons +<br>6 SBCs with two endbulbs and 11 boutons | 13 |
| 40% | 16 SBCs with one endbulb and 11 boutons +<br>4 SBCs with two endbulbs and 10 boutons | 12 |
| 45% | 18 SBCs with one endbulb and 10 boutons +<br>2 SBCs with two endbulbs and 9 boutons | 11 |
| **50%** | 20 SBCs with one endbulb and 9 boutons | **10** |

### Addition of temporal jitter to bushy cell spikes

Temporal disturbance of spike conduction caused by myelinopathy was modeled as added jitter in spike timing of bushy cells. For each bushy cell spike, a random number (in ms) that was drawn independently from an alpha distribution was added to the spike time [26]. The magnitude of jitter (Figs. 5*–*9) was defined as the standard deviation of these alpha-distributed random delays. We changed the scale parameter of the distribution function to obtain resulting jitters in the range of 0.0-1.4 ms.

### Acoustic stimuli

The duration of each repetition of simulation was 500 ms. The AM sound stimulus started at 60 ms and lasted for 420 ms. The first 20 ms of the sound-driven activity was discarded and the remaining 400 ms of response were analyzed. The carrier frequency of the AM sounds and the characteristic frequency of the AN model were both fixed to 7000 Hz. The envelope frequencies used were 150, 300, and 450 Hz (all with 100% modulation depth). The sound intensity was varied in the range of 0-70 dB SPL. For the analysis of binaural tuning, we used 35 dB SPL, at which simulated LSO responses were prominent in comparison to lower or higher levels [8].

### Output measures

Spike rates of the AN fibers, GBCs, SBCs, and LSO were calculated for each 400-ms sound-driven response (see above). Binaural phase-difference tuning of LSO was examined for envelope IPDs ranging between *−*180 and +180 degrees, and the tuning depth was calculated as the difference between maximum spike rate and minimum spike rate. Envelope phase locking was quantified as vector strength [46] and was used for a measure of temporal fidelity in bushy cell spike trains.

### Binaural neuronal discriminability

The neuronal discriminability **D** was calculated for each binaural tuning curve of LSO as in a previous study [23]. In short, the value of **D** at each IPD is defined as the spike rate difference between neighboring IPDs (10 degrees in our simulations) divided by the average standard deviation at these two IPDs. This discriminability measure indicates how well a downstream neuron can theoretically infer the value of IPD by observing the spike rate of the LSO neuron. For ILD-tuning curves, a 5-dB step was used. A three-point smoothing was applied to the resulting curve for **D** to reduce noise that originated from the limited number of repetitions. The average discriminability was calculated as the root mean square of the values of **D** over the range of *±*180 degrees.

## Statistics and Reproducibility

Binaural responses of the LSO model were computed for 100 repetitions (corresponding to 40 seconds of responses in total). Because of the convergence of bushy cell inputs to one LSO neuron (Fig. 1B), 20 and 8 times larger number of repetitions were generated respectively for SBCs and GBCs, each of which in turn required 20 repetitions of AN fiber simulations. Hence 56000 repetitions of AN simulations in total are required for 100 repetitions of LSO. All these repetitions were used for the calculation of output measures (spike rate and vector strength). The bushy cell models as well as the LSO model were deterministic.Therefore, the sources of variability in our series of simulations were the AN spike trains [24] and the added random jitter in bushy cell spike times.

We did not apply any statistical tests for the output measures, because increasing the number of repetitions would lead to statistical significance for any small differences. In the simulation of LSO responses, the mean spike rate may vary by one to a few percent due to trial-to-trial variability for each set of 100 repetitions, which did not affect the overall trends and conclusions. Uncertainty in averaged responses of AN and bushy cells was much smaller because of a larger total number of repetitions.

## Data availability

Simulated data with plotting scripts are available online at [URL will be inserted upon acceptance of the manuscript].

## Code availability

Matlab implementation of the model is openly accessible in the above mentioned data repository.

## Author Contributions

**G.A.:** Conceptualization, Methodology, Software, Formal Analysis, Investigation, Visualization, Writing – Original Draft, Writing – Review and Editing.

**D.J.T.:** Conceptualization, Methodology, Validation, Resources, Funding Acquisition, Writing – Original Draft, Writing – Review and Editing.

